# Cloning of Two Gene Clusters Involved in the Catabolism of 2,4-Dinitrophenol by *Paraburkholderia* sp. Strain KU-46 and Characterization of the Initial DnpAB Enzymes and a Two-Component Monooxygenase DnpC1C2

**DOI:** 10.1101/749879

**Authors:** Taisei Yamamoto, Yaxuan Liu, Nozomi Kohaya, Yoshie Hasegawa, Peter C.K. Lau, Hiroaki Iwaki

## Abstract

Besides an industrial pollutant, 2,4-dinitrophenol (DNP) has been used illegally as a weight loss drug that had claimed human lives. Little is known about the metabolism of DNP, particularly among Gram-negative bacteria. In this study, two non-contiguous genetic loci of *Paraburkholderia* (formerly *Burkholderia*) sp. strain KU-46 genome were identified and four key initial genes (*dnpA*, *dnpB*, and *dnpC1C2*) were characterized to provide molecular and biochemical evidence for the degradation of DNP via the formation of 4-nitrophenol (NP), a pathway that is unique among DNP utilizing bacteria. Reverse transcription PCR analysis indicated that the *dnpA* gene encoding the initial hydride transferase (28 kDa), and the *dnpB* gene encoding a nitrite-eliminating enzyme (33 kDa), are inducible by DNP and the two genes are organized in an operon. Purified DnpA and DnpB from overexpression clones in *Escherichia coli* effected the transformation of DNP to NP via the formation of hydride-Meisenheimer complex of DNP. The function of DnpB appears new since all homologs of DnpB sequences in the protein database are annotated as putative nitrate ABC transporter substrate-binding proteins. The gene cluster responsible for the degradation of DNP after NP formation was designated *dnpC1C2DXFER.* DnpC1 and DnpC2 were functionally characterized as the respective FAD reductase and oxygenase components of the two-component NP monooxygenase. Both NP and 4-nitrocatechol were shown to be substrates, producing hydroquinone and hydroxyquinol, respectively. Elucidation of the *hqdA1A2BCD* gene cluster allows the delineation of the final degradation pathway of hydroquinone to ß-ketoadipate prior to its entry to the tricarboxylic acid cycle.

**Importance:** This study fills a gap in our knowledge and understanding of the genetic basis and biochemical pathway for the degradation of 2,4-dinitrophenol (DNP) in Gram-negative bacteria, represented by the prototypical *Paraburkholderia* sp. strain KU-46 that metabolizes DNP through the formation of 4-nitrophenol, a pathway unseen by other DNP utilizers. The newly cloned genes could serve as DNA probes in biomonitoring as well as finding application in new biocatalyst development to access green chemicals. By and large, knowledge of the diverse strategies used by microorganisms to degrade DNP will contribute to the development of bioremediation solutions since DNP is an industrial pollutant used widely in the chemical industry for the synthesis of pesticides, insecticides, sulfur dyes, wood preservatives, and explosives, etc. (119 words)

## Introduction

2,4-dinitrophenol (DNP) is a yellow crystalline organic compound, industrial uses of which include the production of wood preservatives, sulfur dyes, herbicides, photographic developers and explosives (1, 2). It is listed among the 126 priority pollutants of the United States Environmental Protection Agency regulated in the Clean Water programs (3). In cellular metabolism DNP acts as an ionophore, a classic uncoupler of mitochondrial oxidative phosphorylation made famous in 1961 by Peter Mitchell’s chemiosmotic hypothesis. Interestingly, as early as 1933 the possible therapeutic use of DNP as a weight loss agent in humans was advocated due to its high metabolic rate (4). But unfortunately, serious adverse effects, acute toxicity including deaths due to hyperthermia, for example, had been reported over the years among users (e.g., bodybuilders) of the so-called yellow slimming pill (5). For a historical account and current development see Grundlingh et al. (6); Geisler (7).

In the environment, the major site of DNP degradation is the soil where certain microorganisms can metabolize it. Thus far, only a few bacterial strains capable of growth on DNP as its sole nitrogen or carbon source had been isolated, the majority of which were Gram-positive actinomycetes: *Rhodococcus* and *Nocardioides* (for reviews: 8, 9). Pioneering work of Knackmuss and co-workers who initially used strains of *Rhodococci* (HL 24-1 and 24-2) that were capable of growth on the DNP as well as picric acid (PA; 2,4,6-trinitrophenol) as the sole source of nitrogen and/or carbon, led to the identification of a hydride-Meisenheimer complex of DNP (designated H^−^-DNP; H^−^-PA in the case of PA) as a key intermediate (10). This initial step is the reduction of the aromatic ring, a reaction catalyzed by a NADPH-dependent F_420_ -reductase and hydride transferase system. A second reduction and hydride transferase reaction results in the formation of 2,4-dinitro-cyclohexanone that either goes to the formation of 4,6-dinitrohexanoate or 3-nitroadipate of which the mechanism is not known (Fig. 1A) (11–13). The gene cluster containing the NADPH-dependent F_420_ reductase encoded by *npdG*, and the hydride transferase encoded by *npdI* for the pathway in *R.* (*opacus*) *erythropolis* strain HL PM-1 had been identified (13). Both genes were subsequently found to be highly conserved among several DNP-degrading *Rhodococcus* spp. (13).

**Fig. 1.**
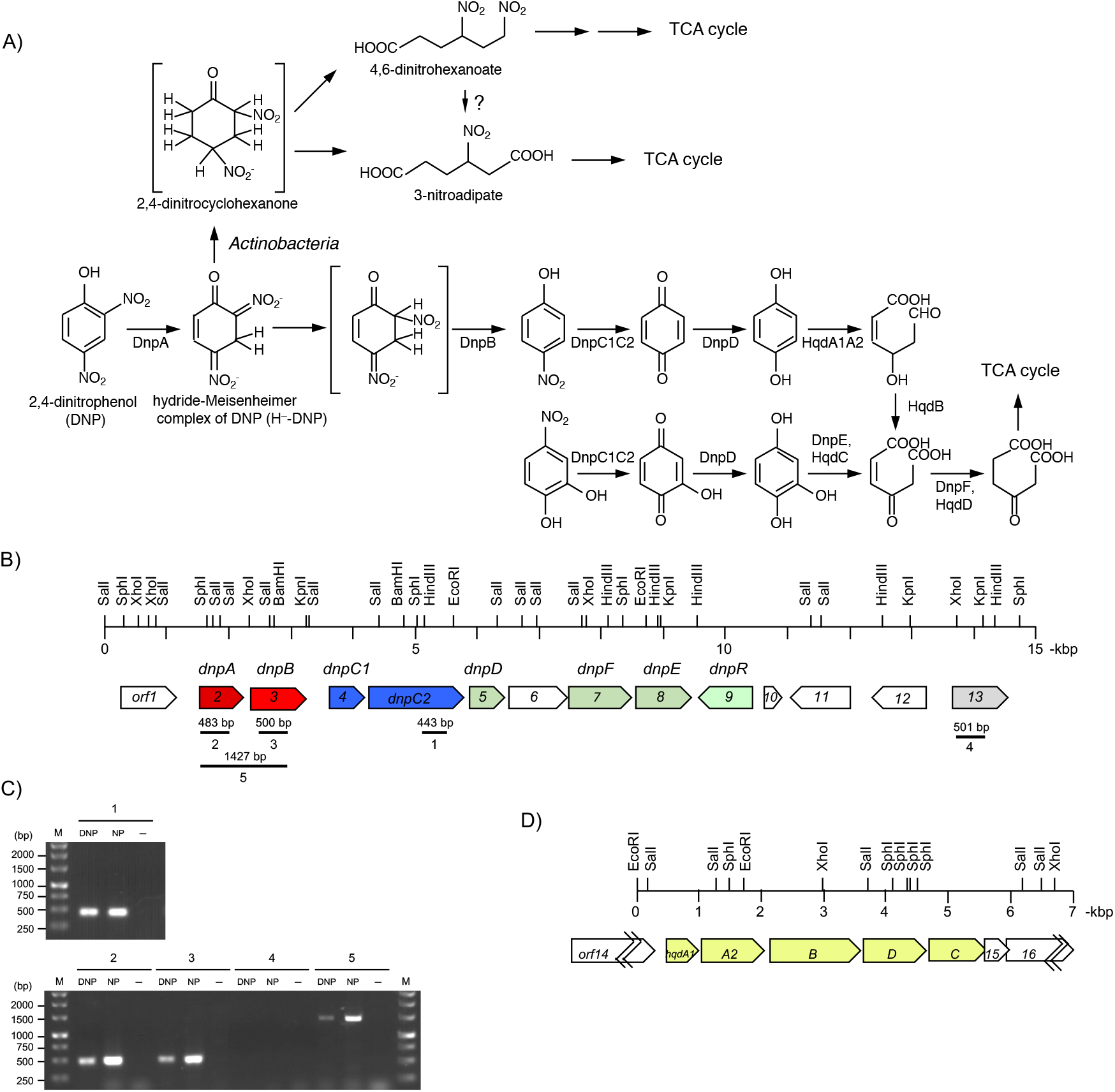
(A) Proposed degradation pathways for 2,4-dinitrophenol (DNP) and 4-nitrocatechol in *Paraburkholderia* sp. strain KU-46, and assignment of the *dnp* gene products to the pathways. For comparison, the commonly known pathway in *Actinobacteria* is shown which does not form nitrophenol as an intermediate. (B) Organization of the *dnp* gene cluster in strain KU-46. See Table 1 for sequence homology searches. The five solid lines (numbered 1 to 5) indicate the size in basepairs of the RT-PCR-amplified regions. (C) Total RNA from strain KU-46 cells grown on MSM medium containing 0.3% succinate and 0.4 mM DNP, NP or NaNO_3_ (-) were used templates. Lane 1, *npdC2* internal region; lane 2, *dnpA* internal region; lane 3, *dnpB* internal region; lane 4, *orf13* internal region; lane 5, *dnpA*-*dnpB* intergenic region. (D) Organization of hydroquinone-degradation gene locus in strain KU-46. *hqdA1, A2, B, C, D* and *orf14*, *15* and *16* are described in Table 1.

**TABLE 1.** BLAST homology search results for deduced amino acid sequences.

| Gene | Most similar gene products (organism)<sup>a</sup> | % identity | score (bits) | E value | Accession no. |
| --- | --- | --- | --- | --- | --- |
| *orf1* | TcpY, Unknown function in 2,4,6-trichlorophenol degradation (*Cupriavidus pinatubonensis* JMP134) | 45.6 | 264 | 8e-84 | AAZ60954 |
| *orf2*<br>(*dnpA*) | WQE_07512, short-chain dehydrogenase/reductase SDR (*Paraburkholderia terrae* BS001) | 99.6 | 492 | 5e-176 | EIN01638 |
| *orf3*<br>(*dnpB*) | WQE_07507, nitrate ABC transporter substrate-binding protein (*Paraburkholderia terrae* BS001) | 99.0 | 577 | 0.0 | EIN01637 |
| *orf4*<br>(*dnpC1*) | HadX, FAD reductase (*Ralstonia pickettii* DTP0602) | 57.8 | 209 | 3e-66 | BAM65765 |
|  | TftC, FAD reductase (*Burkholderia cepacia* AC1100) | 57.8 | 204 | 2e-64 | AAC23547 |
|  | TcpX, FAD reductase (*Ralstonia eutropha* JMP134) | 51.4 | 200 | 2e-62 | AAZ60950 |
| *dnpC2* | TcpA, 2,4,6-trichlorophenol monooxygenase (*Cupriavidus pinatubonensis* JMP134) | 86.1 | 964 | 0.0 | AAM55214 |
|  | HadA, 2,4,6-trichlorophenol monooxygenase (*Ralstonia pickettii* DTP0602) | 85.7 | 955 | 0.0 | BAL41659 |
| *orf5*<br>(*dnpD*) | TcpB, quinone reductase (*Cupriavidus pinatubonensis* JMP134) | 70.3 | 307 | 2e-104 | AAZ60952 |
|  | HadB, probable electron transfer protein (*Ralstonia pickettii* DTP0602) | 68.3 | 282 | 8e-95 | BAM65763 |
| *orf6* | Reut_A1582, Probable FMN adenylyltransferase (*Ralstonia eutropha* JMP134) | 45.6 | 233 | 1e-72 | AAZ60948 |
|  | ORF1, Probable FMN adenylyltransferase (*Ralstonia pickettii* DTP0602) | 48.8 | 233 | 2e-72 | BAL41656 |
| *orf7*<br>(*dnpF*) | TcpD, maleylacetate reductase (*Cupriavidus pinatubonensis* JMP134) | 65.3 | 484 | 4e-169 | AAZ60955 |
|  | HadD, maleylacetate reductase (*Ralstonia pickettii* DTP0602) | 64.5 | 448 | 4e-15 | BAL41322 |
| *orf8*<br>(*dnpE*) | HadC, 6-chlorohydroxyquinol-1,2-dioxygenase (*Ralstonia pickettii* DTP0602) | 69.5 | 402 | 1e-139 | AGW92783 |
|  | TcpC, 6-chlorohydroxyquinol-1,2-dioxygenase (*Cupriavidus pinatubonensis* JMP134) | 68.6 | 398 | 5e-138 | AAM55216 |
| *orf9*<br>(*dnpR*) | HadR, LysR family transcriptional regulator for 2,4,6-trichlorophenol catabolic operon (*Ralstonia pickettii* DTP0602) | 64.4 | 428 | 1e-148 | BAM71407 |
|  | TcpR, LysR family transcriptional regulator for 2,4,6-trichlorophenol catabolic operon (*Cupriavidus pinatubonensis* JMP134) | 65.7 | 417 | 4e-144 | AAZ60949 |
| *orf10* | No significant similarity found |  |  |  |  |
| *orf11* | WQE_07462, AraC family transcriptional regulator (*Paraburkholderia terrae* BS001) | 98.6 | 704 | 0.0 | EIN01628 |
| *orf12* | WQE_07457, LysR family transcriptional regulator (*Paraburkholderia terrae* BS001) | 98.0 | 617 | 0.0 | EIN01627 |
| *orf13* | WQE_07452, NADH:flavin oxidoreductase (*Paraburkholderia terrae* BS001) | 95.1 | 553 | 0.0 | EIN01626 |
| *orf14* | PnpR, LysR-type transcriptional regulator controlling the expression of PNP catabolic operons (*Pseudomonas* sp. WBC-3) | 57.8 | 103 | 1e-24 | AIV98010 |
| *hqdA1* | PnpE2, hydroquinone dioxygenase small subunit (*Burkholderia* sp. SJ98) | 79.8 | 271 | 3e-91 | AFR33818 |
| *hqdA2* | PnpE1, hydroquinone dioxygenase large subunit (*Burkholderia* sp. SJ98) | 92.0 | 635 | 0.0 | AFR33817 |
| *hqdB* | PnpD, 4-hydroxymuconic semialdehyde dehydrogenase (*Burkholderia* sp. SJ98) | 90.1 | 923 | 0.0 | EKS70308 |
| *hqdD* | PnpF, maleylacetate reductase (*Burkholderia* sp. SJ98) | 84.6 | 608 | 0.0 | EKS70307 |
| *hqdC* | PnpC, hydroxyquinol-1,2-dioxygenase (*Burkholderia* sp. SJ98) | 79.3 | 485 | 9e-172 | EKS70306 |
| *orf15* | Orf5, 4-nitrocatechol monooxygenase ferridoxin protein<br>(*Burkholderia* sp. SJ98) | 62.4 | 149 | 2e-44 | EKS70305 |
| *orf16* | SAMN04487926_104331, ABC-2 type transport system<br>ATP-binding protein (*Burkholderia* sp. yr281) | 98.5 | 532 | 0.0 | SDH41711 |

<sup>a</sup>The genes in functionally uncharacterized gene cluster are not considered here, except for *orf1-3*, *orf10-13*, and
*orf16*.

Our laboratory had isolated a Gram-negative *Paraburkholderia* (formerly *Burkholderia*) sp. strain KU-46 that utilized DNP as the sole source of carbon and nitrogen and it degraded DNP via the formation of 4-nitrophenol (NP) and 1,4-benzoquinone (15). A hypothetical reaction mechanism for the formation of NP from DNP was proposed that involved a hydride-Meisenheimer complex, modeled after the formation of H^−^-PA or H^−^-DNP in the degradation pathway of PA/DNP in *Rhodococcus* (9, 10, 16). Interestingly, the cofactor F_420_ system was reported to be limited in its taxonomic distribution and had never been reported in Gram-negative bacteria (17–19). We reckoned that strain KU-46 must likely possess a new enzyme system for the formation of H^−^-DNP. To gain a better understanding of the metabolism of DNP in a Gram-negative bacterium, we carried out molecular analysis and biochemical characterization of some of the key steps in the DNP pathway in strain KU-46. A consolidated pathway including a number of potential genes and regulatory elements is shown in Fig. 1. In addition, we discussed the diversity/evolution of DNP catabolism to related nitroaromatics.

## Results

### Localization of NP monooxygenase encoding gene cluster in strain KU-46

Fig. 1B delineates the genetic locus defined by the following results. First, the nucleotide sequence identity (83.5%) of the 533-bp PCR-amplified product with an equivalent segment of a known oxygenase component of the 2,4,6-trichlorophenol monooxygenase of *Cupriavidus pinatubonensis* JMP134, and 69.6% with the chlorophenol-4-monooxygenase of *Burkholderia cepacia* AC1100 (20, 21) provided an entry point to the genetic information in question. Subsequently, insertional inactivation of the amplified gene by homologous recombination and selection for Km-resistance gave rise to numerous colonies,10 of which were selected and found incapable of growth on NP as a sole nitrogen source. One mutant, designated strain KU-46C2M, the result of a single crossover mutant (Fig. S1) was used for further study. Whereas this mutant was incapable of growth on DNP and NP as a sole carbon source, it was able to grow on DNP as a nitrogen source. Besides, stoichiometric accumulation of NP in the media was observed when DNP in concentration of 0.1 – 1.0 mM was used (Fig. S2). At the lowest concentration, DNP was depleted after 24 h whereas at higher concentrations it took twice as long. Growth of strain KU-46C2M on DNP as a nitrogen source was attributed to the release of one nitro group from DNP as a nitrite. We designated *dnp* for the DNP-degradation pathway genes, and the initially amplified gene as *dnpC2* (Fig. 1B). These results indicated that the *dpnC2* is responsible for DNP degradation and that it is degraded via NP as proposed previously (15).

Flanking regions of the partial *dnpC2* gene were amplified with the aim of localizing the gene cluster for DNP degradation. By primer walking the amplified fragments were sequenced to provide a contiguous segment of 14,808-bp. Computer analysis showed the presence of 14 complete open reading frames (ORFs), 11 of which are on one strand, the remaining in the opposite direction (Fig. 1B).

### Sequence characteristics of *dnpC2* and upstream ORF

The nucleotide sequence of *dnpC2* consists of 1554 bp that is preceded by an appropriate Shine-Dalgarno (SD) sequence, AGGA, 7 bp from the predicted ATG start codon. The 517-residue DnpC2 polypeptide is most similar in sequence to the functionally characterized oxygenase component of the two component NP and chlorophenol monooxygenases, viz., 2,4,6-trichlorophenol monooxygenase TcpA from *Cupriavidus pinatubonensis* JMP134 (86.1% identity) (21); 2,4,6-trichlorophenol monooxygenase HapD from *Ralstonia pickettii* DTP0602 (85.7% identity) (22); and chlorophenol-4-monooxygenase TftD from *Burkholderia cepacia* AC1100 (65.9% identity) (20).

Upstream of *dnpC2* and arranged in the same direction is *orf4* that consists of 193 codons, and preceded by a consensus SD sequence, GGAG. The translated product of *orf4* resembles the FAD reductase component of the two-component phenol monooxygenases: e.g., HadX of 2,4,6-trichlorophenol monooxygenase from *Ralstonia pickettii* DTP0602 (57.8%) (22), TftC of chlorophenol-4-monooxygenase from *Burkholderia cepacia* AC1100 (57.8%) (20), and TcpX of 2,4,6-trichlorophenol monooxygenase from *Cupriavidus pinatubonensis* JMP134 (51.4% identity) (21). On the basis of the results of homology searches, *orf4* is designated *dnpC1* and predicted to encode the reductase component of the two-component NP monooxygenase system, evidence of which is presented below.

### Functional analysis of *dnpC1* and *dnpC2* genes

At first, overexpression clones of DnpC1C2 and the two individually were verified by SDS-PAGE for their protein production (Fig. S3). The molecular mass of His_10_-tagged DnpC1 (H_10_-DnpC1) was determined to be 24 kDa in good agreement with the predicted relative molecular mass (*M*_r_) of 22,441. Those of DnpC2 and His_10_-tagged DnpC2 (H_10_-DnpC2) were 58 kDa and 59 kDa, respectively, the corresponding predicted *M*_r_ being 58,339 and 60,266, respectively.

*E. coli* whole cells harboring *dnpC1C2* in pET-dnpC1C2 expression plasmid converted NP to hydroquinone at a rate of 0.43 mM·hr^-1^ (Fig. 2, Fig. S4). At a cell density of 1.0 at OD_600_ these cells converted 0.1 mM NP completely within 20 min. On the other hand, *dnpC1* (pET-dnpC1) alone was not able to transform NP, but *dnpC2* (pET-dnpC2) gave a detectable amount of hydroquinone in the reaction mixture (Fig. 2, Fig. S4). The conversion rate was 22% compared to that of the full gene complement, *dnpC1C2*. 4-Nitrocatechol was also transformed at a rate of 0.87 mM·hr^-1^ by DnpC1C2 giving hydroxyquinol as a product (Fig. 2, Fig. S4).

**Fig. 2.**
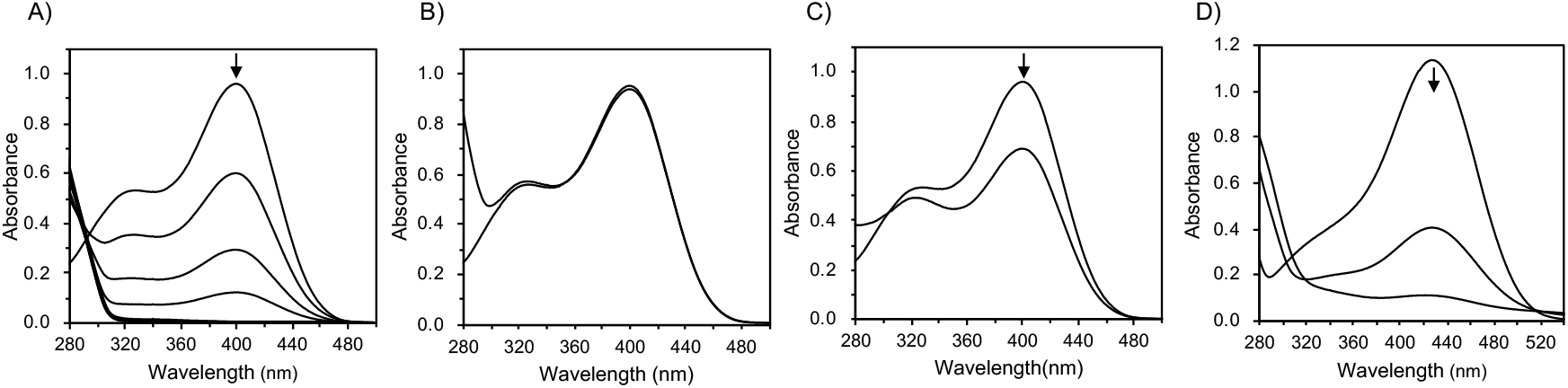
Functional analysis of *dnpC1* and *dnpC2* genes. A) Spectrophotometric changes during the transformation of NP by whole cells of *E. coli* harboring *dnpC1C2* (pET-dnpC1C2). Decrease in spectral absorption at 400 nm correspond to depletion of NP in a time dependent manner. Spectra were recorded every 5 min. B) Spectrophotometric change was not observed during the transformation of NP by whole cells of *E. coli* harboring *dnpC1* (pET-dnpC1), indicating DnpC1 lacked NP transformation activity. Spectra were recorded at 0 and 30 min. C) Spectrophotometric change during the transformation of NP by whole cells of *E. coli* harboring *dnpC2* (pET-dnpC2), indicating partial transformation of NP by DnpC2. Spectra were recorded at 0 and 30 min. D) Spectrophotometric changes during the transformation of 4-nitrocatechol by whole cells of *E. coli* harboring *dnpC1C2* (pET-dnpC1C2). Decrease in spectral absorption at 426 nm corresponds to depletion of 4-nitrocatechol in a time dependent manner. Spectra were recorded every 5 min.

### Additional NP and 4-nitrocatechol degradation genes and unknown ORFs

Downstream of *dnpC2* and on the same DNA strand are *orf5, 6, 7*, and *8*, followed by *orf9* on the opposite strand (Fig. 1B). On the basis of the sequence similarities in addition to the analogy with the 2,4,6-trichlorophenol degradation pathway of *Cupriavidus pinatubonensis* JMP134 (23) and *Ralstonia pickettii* DTP0602 (22), we designated *orf5, 7, 8,* and *9* to the DNP and NP-degrading genes, *dnpD, dnpF, dnpE* and *dnpR*, respectively (Table 1, Fig.1, Fig. S5). The *dnpD, dnpF, dnpE* and *dnpR* genes and *orf6* were predicted to encode quinone reductase, maleylacetate reductase, hydroxyquinol-1,2-dioxygenase, the LysR family transcriptional regulator and FMN adenylyltransferase, respectively.

The *orf10* product has no apparent sequence homolog in the protein database whereas *orf11* and *orf12* products showed significant sequence identity to AraC family transcriptional regulator and LysR family transcriptional regulator, respectively. The *orf13* product showed similarity with putative oxidoreductases that contain the old yellow enzyme (OYE)-like FMN binding domain. Some of the OYE family enzymes have been found to catalyze formation of hydride–Meisenheimer complex from 2,4,6-trinitrotoluene (24–26). However, the C-terminus region of Orf13 appeared to be truncated by about 90 amino acids compared to the homologs.

### Establishment of the initial genes responsible for the conversion of DNP to NP and functional analysis

Upstream of *dnpC1* (*orf4*) and in the same direction, there are three ORFs within the sequenced region by which we employed RT-PCR method to test if they are inducible and the designate for the initial genes of the DNP degradation pathway using *orf2* and *orf13* as candidates, and *dnpC2* as a positive control of NP and DNP induced genes. Amplification of a 1.4-kb fragment indicated that *orf2* and *orf3* are on an operonic unit as suggested by the proximity of the two genes (Fig. 1C). We designated *orf2* and *orf3* as *dnpA* and *dnpB*, respectivety. On the other hand, *orf13* is not induced by DNP or NP.

Characteristics of *dnpA* and *dnpB* are as follows: The coding sequence of *dnpA* consists of 738 nucleotides with an appropriately positioned consensus SD sequence, GGAGGT, 7 bp from the putative ATG start site. DnpA shares 99.6% sequence identity with a hypothetical short-chain dehydrogenase/reductase (SDR) of an unknown function from *Paraburkholderia terrae* BS001 (Table 1). The most similar protein in Protein Data Bank (PDB) is 3-oxoacyl-(acyl-carrier-protein) reductase (FabG1) from *Staphylococcus aureus* subsp. aureus NCTC 8325 (PDB acc. no. 3SJ7) (27) with 36% identity. A predicted secondary structure of DnpA is shown in Fig. S6. The secondary structure elements are similar to those of FabG1 (Fig. S6). Notable sequence features of DnpA are the conservation of a cofactor-binding motif, TGxxxGIG, near the N-terminus of the protein (28) and in addition two key Arg residues which are characteristics of NADP(H)-preferred enzymes (Fig. S6). DnpA sequence, however, lacks a conserved tyrosine (replaced by Leu) in the catalytic YxxxK motif of known SDRs (28).

The *dnpB* gene is located downstream of *dnpA* in the same direction, and separated by a 90-bp intergenic sequence that is preceded by a consensus SD sequence (GAGGT); it is 867-bp and encodes a polypeptide of 288 residues. The BLAST search revealed homologous sequences that were annotated as putative ABC transporter substrate-binding proteins. Specifically nitrate ABC transporter substrate-binding protein of *Paraburkholderia terrae* BS001 scored the highest (Table 1). In terms of Conserved Protein Domain search, DnpB falls in the family of periplasmic-binding protein type2-superfamily.

Transformation assays with whole cells of *E. coli* harboring pET-dnpAB, which expresses DnpA and DnpB, measured by UV-visible light absorption spectrum indicated time-dependent shifts of absorption maxima at 360 nm to 400 nm. This corresponded to the depletion of DNP and the production of NP (Fig. 3A) identified by HPLC with a single peak at a retention time of 3.7 min and a photodiode array spectrum (Fig. S7), indicating that *dnpAB* are the initial genes responsible for the degradation of DNP. We also performed transformation assay of PA (a non-growth substrate of strain KU-46) with whole cells of *E. coli* harboring pET-dnpAB. Time-dependent shifts of absorption maxima at 355 nm to 400 nm corresponding to the depletion of PA was observed with the production of NP identified by HPLC with a single peak at a retention time of 3.9 min (Fig. 4, Fig. S8).

**Fig. 3.**
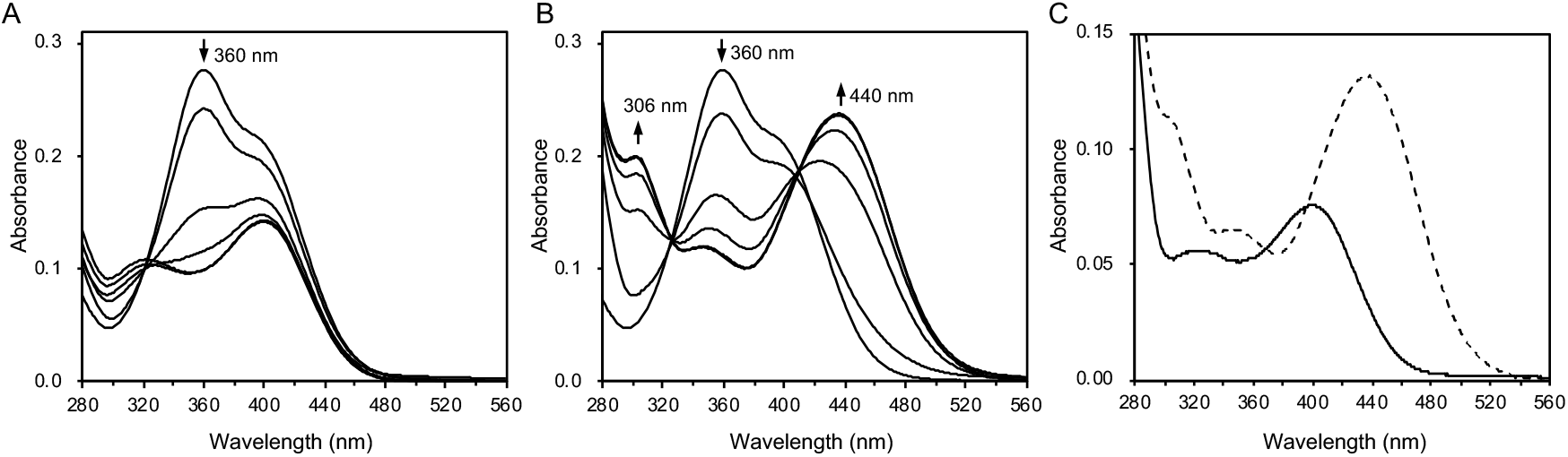
Functional analysis of *dnpAB* genes. A) Spectrophotometric changes during the transformation of DNP by whole cells of *E. coli* harboring *dnpAB* (pET-dnpAB). Spectra were recorded before the addition of cells and after 10, 20, 30 and 40 min. B) Spectrophotometric change during the transformation of DNP by whole cells of *E. coli* harboring *dnpA* alone (pET-dnpA). Spectra were recorded before the addition of cells and after 1, 2, and 3 hrs. The arrows indicate the direction of spectral changes. C) Spectrophotometric change during the transformation of H^−^-DNP by whole cells of *E. coli* harboring *dnpB* alone (pET-dnpB). Dashed line, initial spectrum of H^−^-DNP; the solid line, the final spectrum of the transformation product. Dashed line, initial spectrum of DNP; the solid line, the final spectrum of the transformation product.

**Fig. 4.**
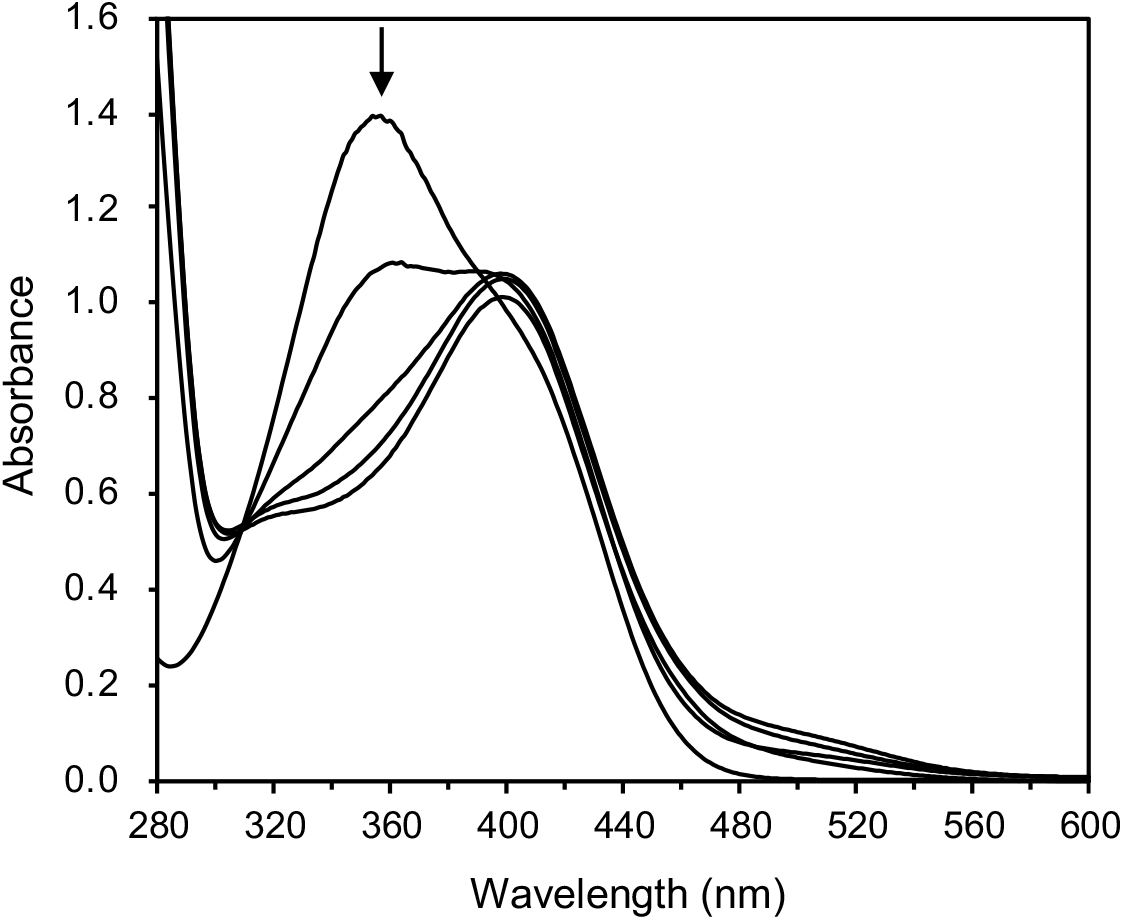
Spectrophotometric changes during the transformation of PA by whole cells of *E. coli* harboring *dnpAB* (pET-dnpAB). The arrows indicate the direction of spectral changes. Spectra were recorded before the addition of cells and after 1, 2, 3 and 4 hrs.

To analyze the function of DnpA and DnpB, His-tagged proteins designated H_10_-DnpA and H_10_-DnpB were produced in *E. coli* containing plasmids pET-dnpA and pET-dnpB, respectively. Analysis of the purified proteins on SDS-PAGE correctly verified the expected molecular masses of the two proteins (Fig. S9). The experimental and predicted mass of H_10_-DnpA are 32 and 28.121; those of H_10_-DnpB are 30 and 33.465 kDa. In the conversion of DNP by whole cells of *E. coli* harboring pET-dnpA, or *dnpA* alone, the transformation product was identified as H^−^-DNP with the characteristic UV-visible light absorption spectrum, λ_max_ =306 and 440 (Fig. 3B) (29). Appearance of an isosbestic point at 405 nm indicated that DNP was transformed to H^−^-DNP (Fig. 3B) (29). This was supported by HPLC analysis showing an identical retention time (2.6 min), and UV-visible spectrum with photodiode array detector, λ_max_ =258, 306, and 440, of the product with the synthetic H^−^-DNP (Fig. S10). On the other hand, *E. coli* whole cells harboring pET-dnpB, or *dnpB* alone, caused a hypsochromic shift of absorption maxima at 430 nm to 400 nm in the conversion of H^−^-DNP (Fig. 3C). This corresponded to depletion of H^−^-DNP and the production of NP. Supporting data for this came from HPLC analysis that showed an identical retention time (3.7 min), and UV-visible spectrum with photodiode array detector, λ_max_ =400 nm, of the product with authentic NP (Fig. S11). The overall results indicated that *dnpA* encodes a hydride transferase catalyzing the conversion of DNP to H^−^-DNP, and *dnpB* encodes a nitrite-eliminating enzyme that transforms H^−^-DNP to NP and nitrite.

We also performed transformation assay of PA by whole cells of *E. coli* harboring *dnpA* alone. In the conversion of PA by the cells, a bathochromic shift was detected, and the transformation product was identified as 2H^−^-PA with the characteristic UV-visible light absorption spectrum λ_max_ =390, and a shoulder at 440 to 480 nm (Fig. S12A) (13). 2H^−^-PA was also converted to NP by *E. coli* whole cells harboring *dnpB* alone (Fig. S12B).

### Transformations by purified DnpA and DnpB

To exclude any action of *E. coli* inherent enzyme in the transformation of DNP and H^−^-DNP, and to analyze possible coenzyme requirement, transformation assays were performed with purified HAT-tagged DnpA (H-DnpA) and H_10_-DnpB (Fig. S9). Transformation of DNP to H^−^-DNP with purified H-DnpA was achieved when NADPH was used in the reaction mixture (Fig. 5; Fig. S13), whereas NADH did not work. On the other hand, transformation of H^−^-DNP to NP with purified H_10_-DnpB did not require any cofactor (Fig. 6; Fig. S14).

**Fig. 5.**
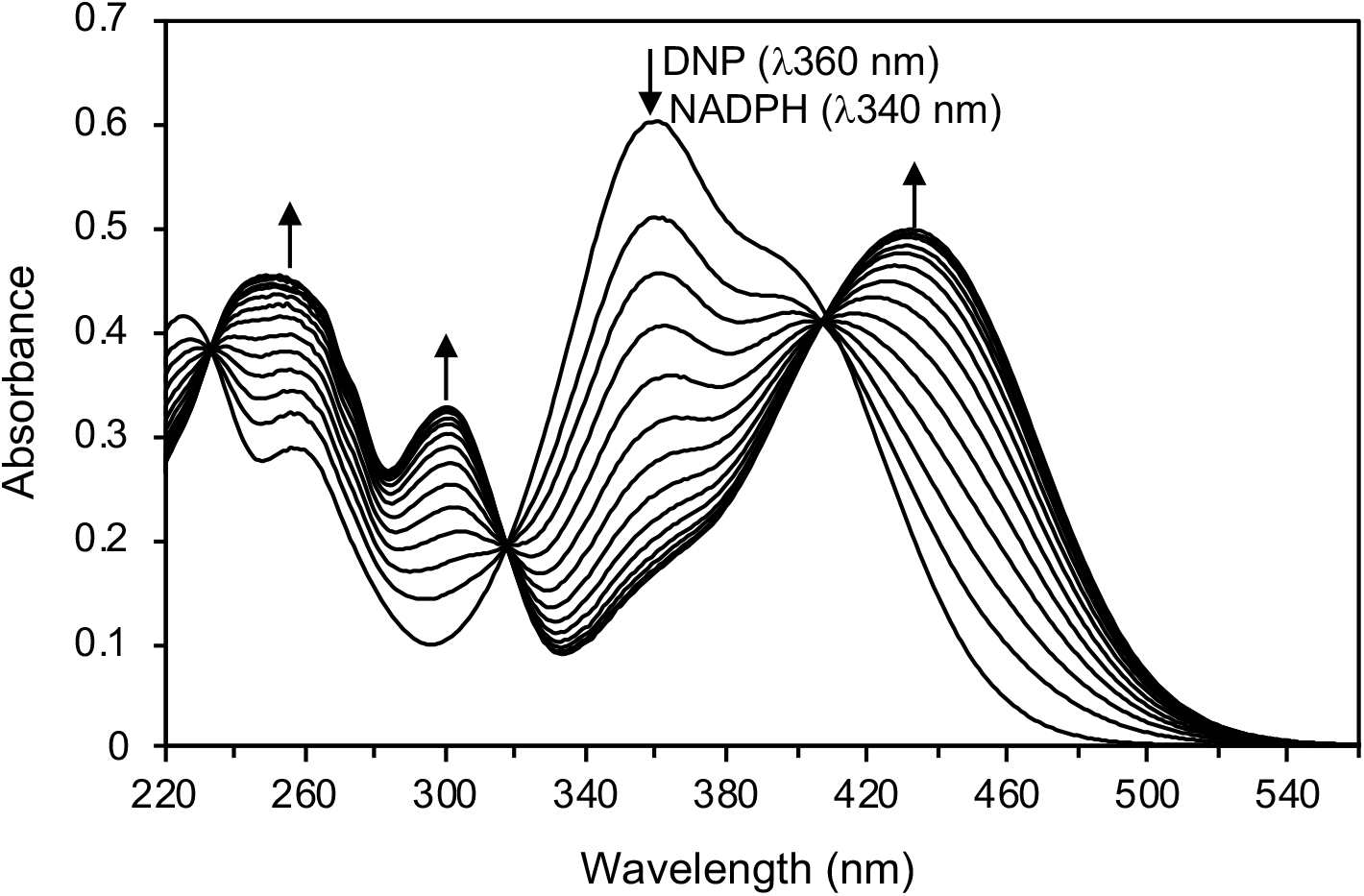
Spectrophotometric change during the transformation of DNP by purified H-DnpA. Sample and reference cuvettes contained NADPH (0.08 μmol), sodium-potassium phosphate buffer (20 μmol, pH 7.1), and 1 μg H-DnpA in a 1 ml volume. The sample cuvette also contained 0.04 μmol of DNP. The arrows indicate the direction of spectral changes.

**Fig. 6.**
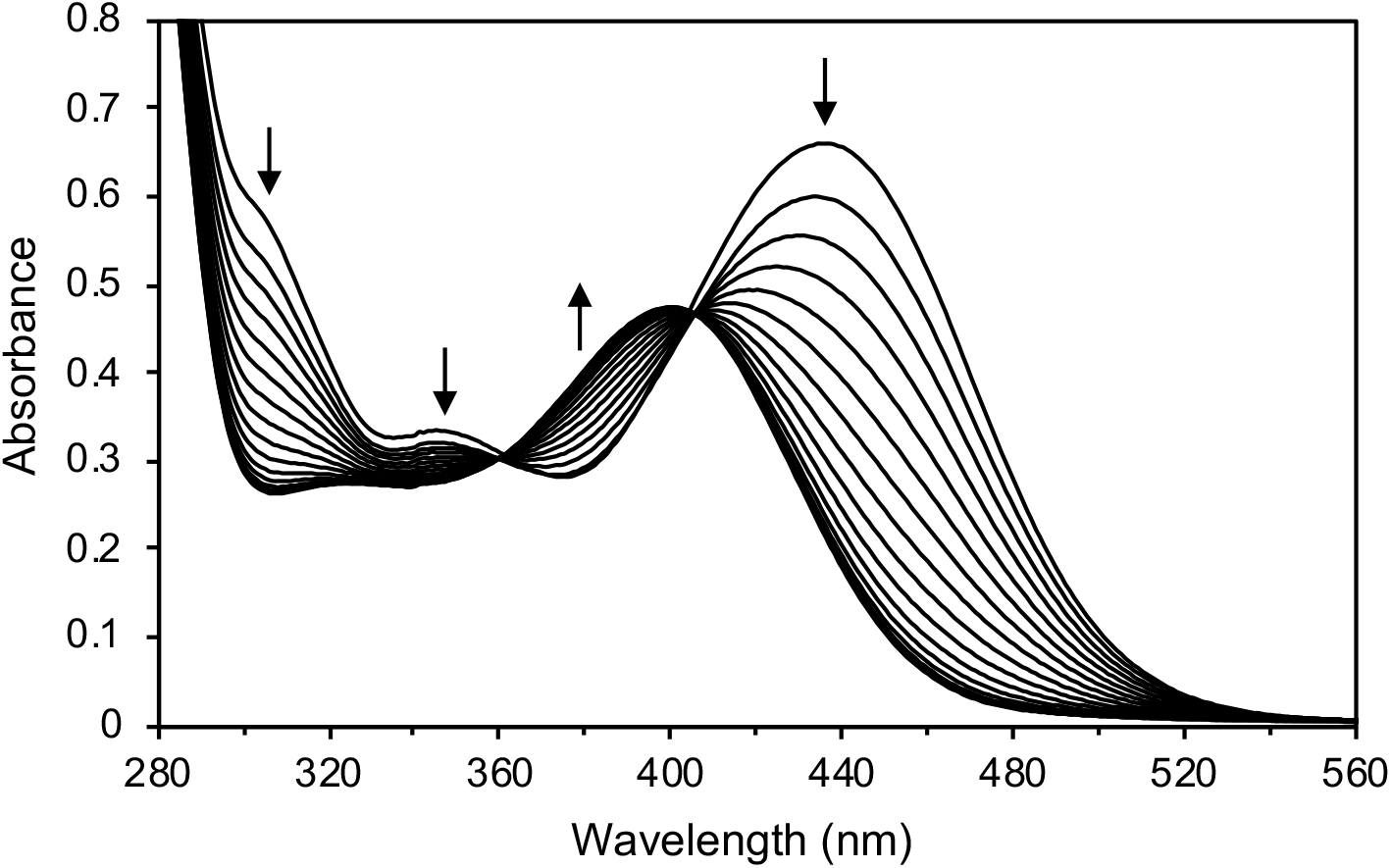
Spectrophotometric change during the transformation of H^-^-DNP by purified H_10_-DnpB. Sample and reference cuvettes contained sodium-potassium phosphate buffer (20 μmol, pH 7.1), and 1.0 μg H_10_-DnpB in a 1 ml volume. The sample cuvette also contained 0.05 μmol of H^-^-DNP. The arrows indicate the direction of spectral changes.

We also performed transformation assays of non-growth substrate of strain KU-46, PA, with purified H-DnpA and H_10_-DnpB. In the transformation of PA by H-DnpA, a two-step time-dependent shifts of absorption maxima of UV-visible light absorption spectra were observed (Fig. 7A). In the first step, a bathochromic shift was detected (Fig. 7B). The transformation product was identified as hydride Meisenheimer complex of PA (H^−^-PA) with the characteristic UV-visible light absorption spectrum features, λ_max_=420 nm and a shoulder at around 490 nm (13, 30). In the second step, a hypsochromic shift from λ_max_=420 to 400 indicated H^−^-PA was transformed to dihydride Meisenheimer complex of PA (2H^−^-PA) (Fig. 7C) (13). The formation of both H^−^-PA and 2H^−^-PA was supported by HPLC analysis (Fig. S15A-C). After the purified H-DpnA was removed from the reaction mixture by Amicon Ultra-4 10 kDa (Merck Millipore), purified H_10_-DnpB was added to the reaction mixture. Time-dependent shifts of absorption maxima of UV-visible light absorption spectra with two steps were also observed involving at first (Fig. 7D). A bathochromic shift and a formation of second absorption maxima at 306 nm, indicative of formation of H^−^-DNP (Fig. 7E) followed by a hypsochromic shift to λ_max_=400, indicating formation of NP (Fig. 7F). The formation of NP was supported by HPLC analysis (Fig. S15D). HPLC analysis also showed the formation of DNP (Fig. S15D).

**Fig. 7.**
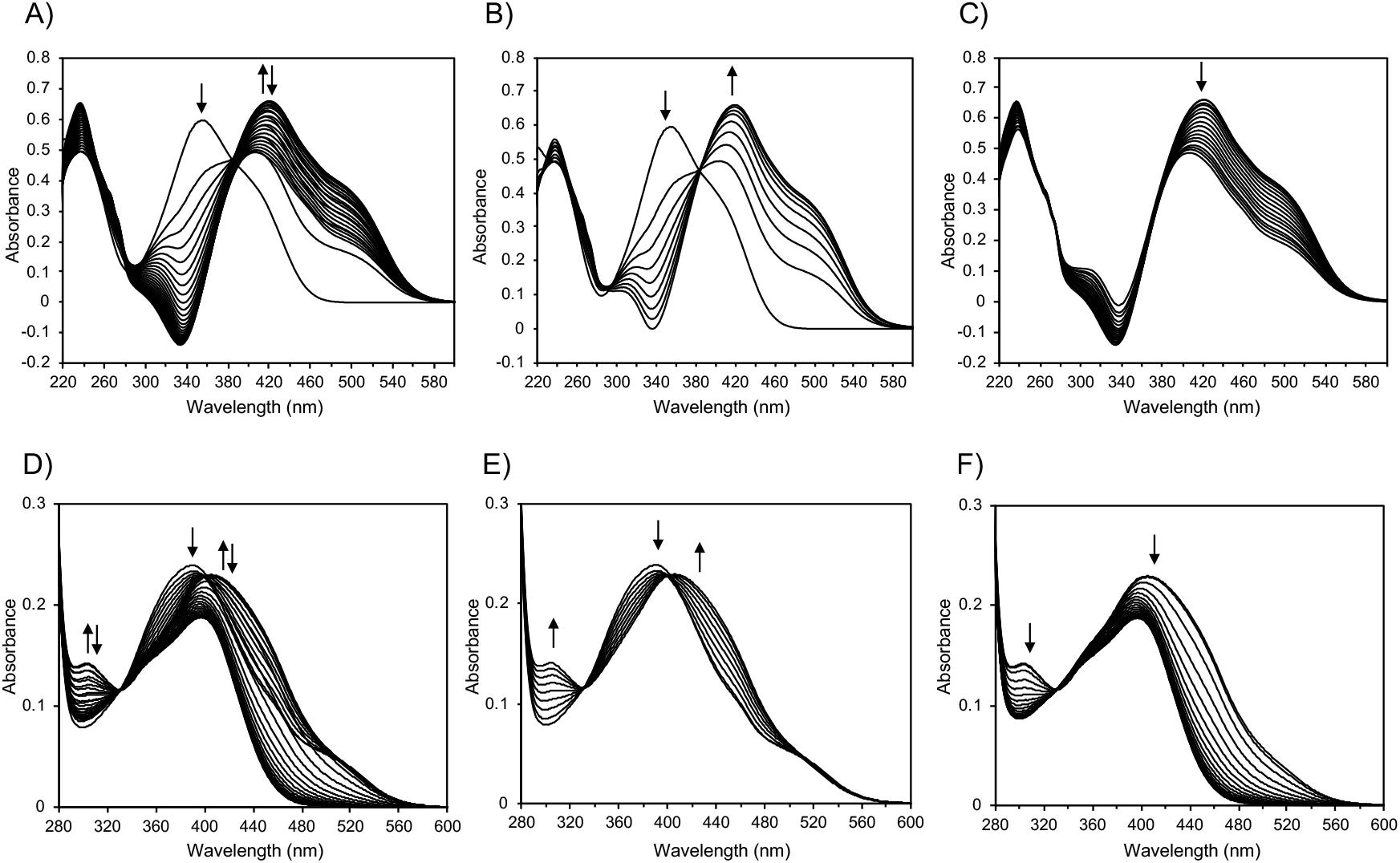
Spectrophotometric changes during the transformation of PA by purified H-DnpA and H_10_-DnpB. The spectra were recorded every 2 minutes. The arrows indicate the direction of spectral changes. A) Spectrophotometric change during the transformation of PA by purified H-DnpA. Sample and reference cuvettes contained NADPH (0.08 μmol), sodium-potassium phosphate buffer (20 μmol, pH 7.1), and 1 μg H-DnpA in a 1 ml volume. The sample cuvette also contained 0.04 μmol of PA. The arrows indicate the direction of spectral changes. Time-dependent shifts of absorption maxima of UV-visible light absorption spectra with two steps were observed. The shift of absorption maxima from 355 nm to 420 nm was observed from 0 to 18 min, and the shift of absorption maxima from 420 nm to 400 nm was observed after 18min. B) Spectrophotometric change from 0 to 18 min of panel A. C) Spectrophotometric change after 18 min of panel A. D) Spectrophotometric change during the transformation of the product of panel A by purified H_10_-DnpB. Time-dependent shifts of absorption maxima of UV-visible light absorption spectra with two steps were observed. A bathochromic shift and a formation of second absorption maxima at 306 nm were observed from 0 to 16 min, and decrease in spectral absorption at 306 nm and a hypsochromic shift were observed after 16 min. E) Spectrophotometric change from 0 to 16 min of panel D. F) Spectrophotometric change after 16 min of panel D. Spectrophotometric change from 0 to 16 min of panel D.

### Localization of hydroquinone degradation genes

The genes encoding hydroquinone dioxygenase and 4-hydroxymuconic semialdehyde dehydrogenase were absent in the sequenced 15-kb region. Amplification by the degenerate primer sets (hqdA2-F and hqdA2-R, and hqdB-F and hqdB-R; Table 2), gave rise to DNA fragments of 686-bp and 986-bp, respectively. As expected, the deduced amino acid sequences from these fragments showed similarity to those of known hydroquinone dioxygenase large subunit, and 4-hydroxymuconic semialdehyde dehydrogenase, respectively (Table 1). Furthermore, using primer set hqdA2-F and hqdB-R, a 2.2-kb DNA fragment was obtained, indicating *hqdA2* and *hqdB* are adjacent to each other in the order of *hqdA2*-*hqdB*. The flanking regions of the 2,208-bp region were amplified to isolate the hydroquinone degradation genes. Consequently, from a contiguous segment of sequenced DNA of 6,754-bp, six complete ORFs and 2 partial ORFs were obtained (Fig. 1D). On the basis of their sequence relatedness to known proteins ranging from 80-92% identity (Table 1), the first two ORFs were designated *hqdA1* and *hqdA2,* to encode the small subunit and large subunit of hydroquinone dioxygenase, respectively. The *hqdB, hqdD* and *hqdC* genes were predicted to encode hydroxymuconic semialdehyde dehydrogenase, maleylacetate reductase, and hydroxyquinol-1,2-dioxygenase, respectively.

**TABLE 2.**
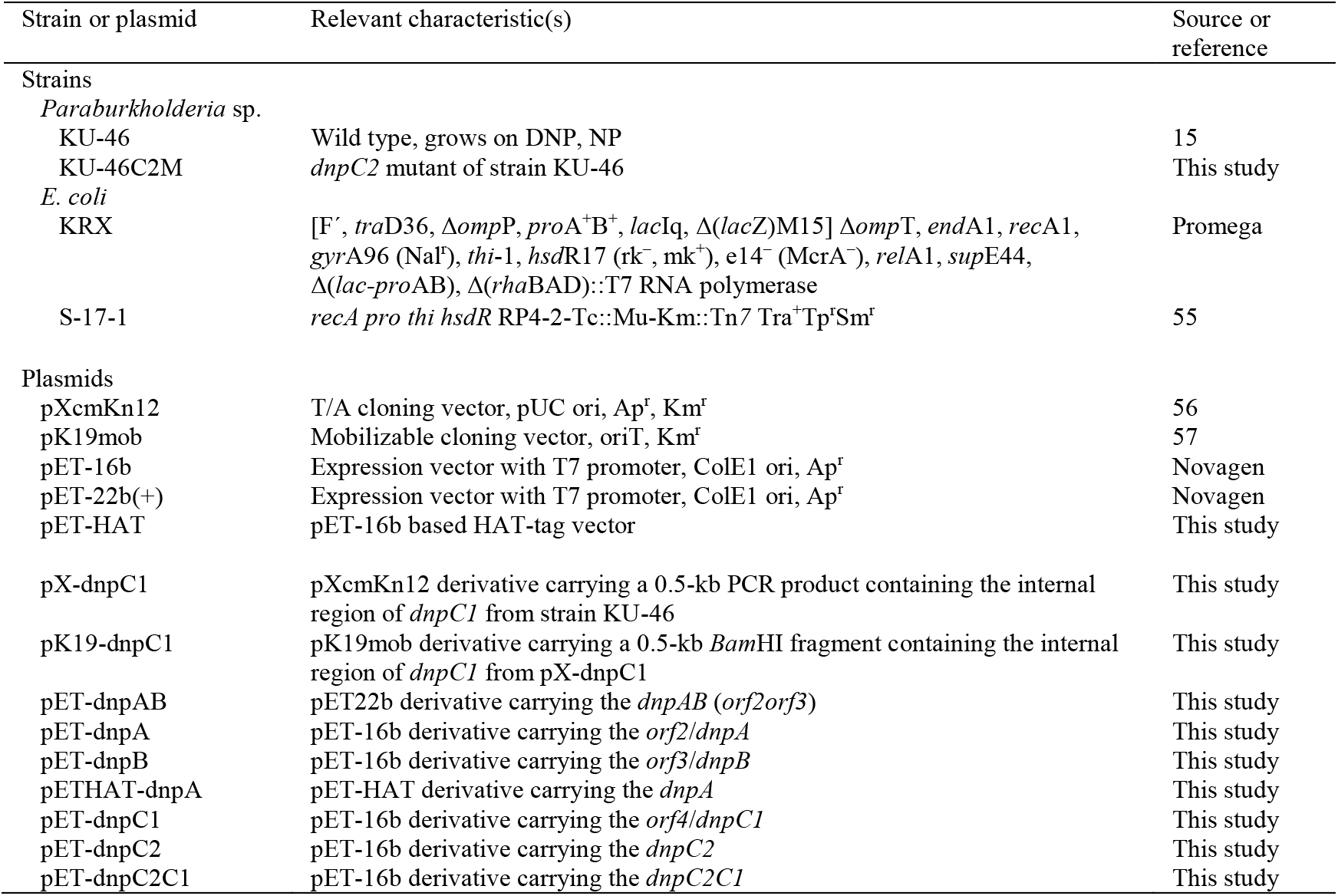
Bacterial strains and plasmids used in this study.

## Discussion

In this study, a molecular view of DNP metabolism by a Gram-negative bacterium is surfacing for the first time. The genetic basis for the initial reaction, namely the removal of a nitrite from DNP to form NP via a hydride-Meisenheimer complex, is attributed to the co-transcribed *dnpA* and *dnpB* genes that encode a 28-kDa hydride transferase (DnpA) and a 33-kDa nitrite-eliminating enzyme (DnpB), respectively (Fig. 1). Both DnpA and DnpB are new entities. DnpA is a novel member of the large superfamily of SDRs since no such ascribed activity had been reported previously (31). Interestingly, like DnpA, of which a oxoacyl-(ACP) reductase is the closest homolog, that of ANI02794.1, was recently found to function as a 17β-hydroxysteroid dehydrogenase in the conversion of 17β-estradiol into estrone in *Pseudomonas putida* SJTE1 (32). Members of SDRs are known to have diverse functions and they are distantly related with typically 20-30% residue identity in pair-wise comparisons. Structurally, DnpA was predicted to share similar features of FabG1 from *S. aureus* (PDB:3SJ7), an enzyme that utilizes NADPH to reduce β-ketoacyl-ACP to (*S*)-β-hydroxyacyl-ACP. As expected of the conserved sequence motifs in DnpA (Fig. S6) the transformation of DNP to H^−^-DNP with purified DnpA was achieved in the presence of NADPH. On the other hand, transformation of H^−^-DNP to NP with purified DnpB did not require any of the nicotinamide cofactor. DnpB belongs to the type 2 periplasmic binding fold superfamily, the majority of which are involved in the uptake of a variety of soluble substrates such as phosphate, sulfate, nitrate, polysaccharides, lysine/arginine/ornithine, and histidine (33). However, this family also includes ionotropic glutamate receptors and unorthodox sensor proteins involved in signal transduction. Hence, it should come to no surprise that one such member would have a catalytic activity, two previous examples being 2’-hydroxybiphenyl-2-sulfinate desulfinase (DszB) (34), and THI5: amino-5-hydroxymethyl-2-methylpyrimidine phosphate synthase (THI-5) (35).

Interestingly, DnpAB also transformed PA to NP and DNP, despite PA was not utilized by strain KU-46 as a growth substrate. We proposed the reaction sequence of NP and DNP formation from PA as shown in Fig. 8. Our results indicated that DnpA catalyzes the two sequential reactions: PA to H^−^-PA and H^−^-PA to 2H^−^-PA. In Gram-positive actinomycetes, the formation of 2H^−^-PA is catalyzed by two enzymes – first, a hydride transferase II encoded by *npdI* catalyzes the formation of H^−^-PA from PA, and second, a hydride transferase I encoded by *npdC* catalyzes the formation of 2H^−^-PA from H^−^-PA (36). The formation of DNP and NP from a mixture of 2H^−^-PA and H^−^-PA during DnpB reaction without DnpA indicated that DnpB catalyzes nitrite-elimination from both H^−^-PA and 2H^−^-PA. In contrast, nitrite-eliminating enzyme from Gram-positive actinomycetes was assumed to only accept the 2H^−^-PA as substrate due to the fact that DNP was not produced from PA in Gram-positive actinomycetes. Despite that DnpAB catalyzed the transformation of PA to NP, PA was not utilized as nitrogen and/or carbon source for growth of strain KU-46. This suggests that transcriptional regulator for *dnpAB* does not recognize PA as inducer, i.e., transcription of *dnpAB* is not activated by PA.

**Fig. 8.**
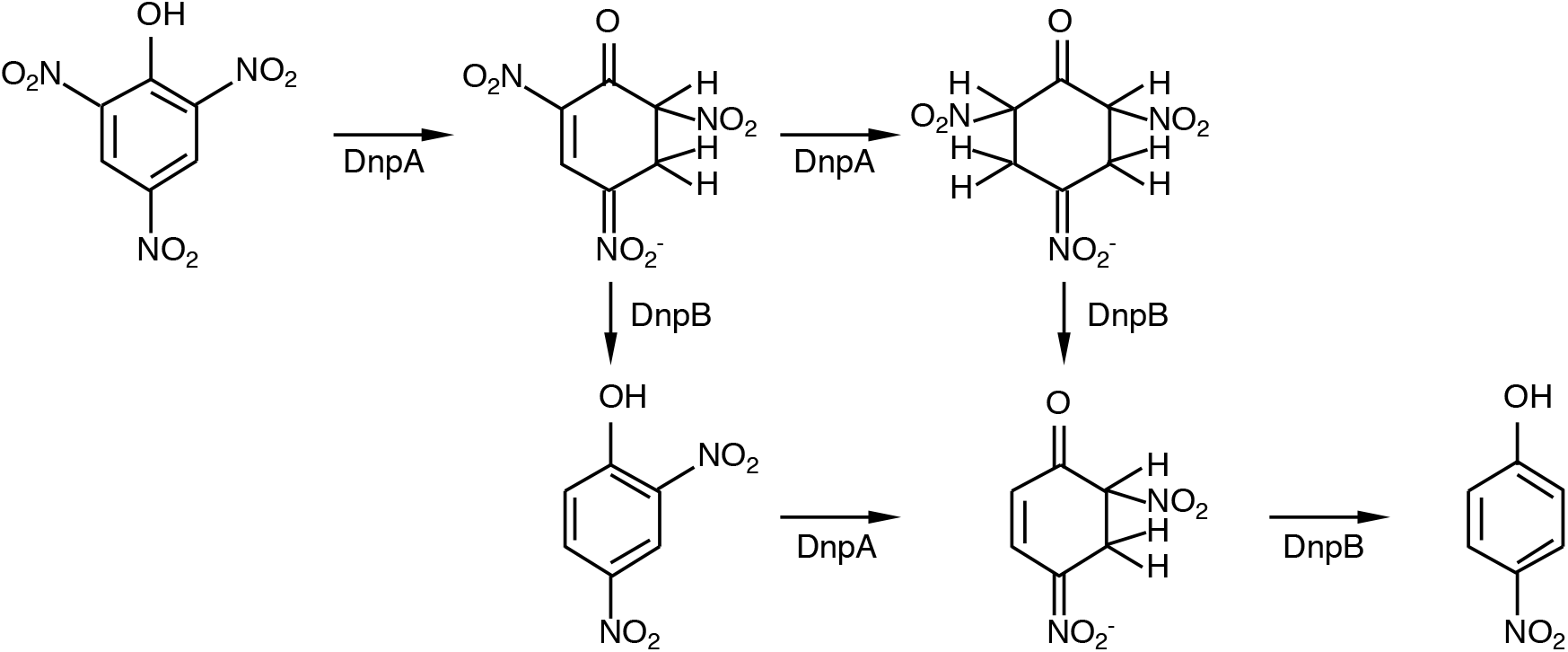
Proposed reaction sequence of NP and DNP formation from PA by DnpA and DnpB.

From a genomics perspective, both *dnpA* and *dnpB* genes and their organization are conserved in the various completely sequenced genomes of *Paraburkholderia* spp. (Fig. S16, Table S1). This mirrors the conserved nature of the *npdG* and *npdI* genes among numerous *Rhodococci* capable of degrading DNP and NP where the 27 kDa NADPH-dependent F_420_ reductase and the 32.9 kDa hydride transferase (HTII), unrelated in sequence or structure to DnpA and DnpB, respectively, had evolved to carry out the same nitrite removal via the formation of hydride Meisenheimer complexes (13, 14). The NpdG functions to shuttle the hydride ions from NADPH to F_420_, the biosynthesis of which is apparently absent in Gram-negative bacteria but instead where the redox factors FAD, FMN and NAD(P) are ubiquitous (18).

Initial access to the *dnp* genetic locus of *Paraburkholderia* sp. strain KU-46 was made possible by homology probing that led to the identification of the two-component *dnpC1C2* genes encoding the reductase and oxygenase components of the NP monooxygenase. This appears to be the first report for a two-component NP monooxygenase from the genera *Burkholderia* and *Paraburkholderia* while a single component NP monooxygenase (PnpA) was described recently in *Burkholderia* sp. SJ98 for the degradation of 3-methyl-4-nitrophenol to methyl-1,4-benzoquinone as the first intermediate (37). However, several two-component *para*-nitrophenol monooxygenases have been reported in Gram-positive bacteria such as *Rhodococcus imtechensis* RKJ300 that initiates the degradation of 2-chloro-4-nitrophenol (38) or the *hadXA* genes of *Ralstonia pickettii* DTP0602 involved in the degradation of halogenated phenols and nitrophenols (39), among many others (23, 40-43; for reviews: 8, 9, 44). Of particular interests are the HadX reductase of *R. pickettii* DTP0602 and TcpA oxygenase components of 2,4,6-trichlorophenol monooxygenase of *C. pinatubonensis* JMP134, which are most identical in sequence to the respective DnpC1 and DnpC2 (Table 1). By and large, the 2,4,6-trichlorophenol monoxygenases of both the TcpXA and HadXA systems are most related to DnpC1C2 and the homologies extend beyond to include putative quinone reductase (DnpD), maleylacetate reductase (DnpF), hydroxyquinol-1,2-dioxygenase (DnpE), and the LysR-type transcriptional regulator (DnpR) (Fig. S5). Evidently, some gene rearrangements had occurred among the three organisms. In at least the sequenced genome of strain DTP0602 it is known that the *hadRXABC* gene cluster is separated from that of *hadSYD* by 146-kb where *hadD* is the maleylacetate reductase encoding gene (22, 45). Whereas in strain KU-46, this same gene (*dnpF*) is only 0.9-kb downstream of *dnpD* (quinone reductase), a similar situation found in strain JMP134 (Fig. S5).

In the biotransformation, *E. coli* whole cells harboring *dnpC1C2* converted NP to hydroquinone, and 4-nitrocatechol to hydroxyquinol. Benzoquinone was not detected in the *E. coli* whole cell transformation assay probably due to the action of an unknown broad substrate reductase(s) in *E. coli* (46, 47). In strain KU-46, the responsible reductase would be DnpD, a protein yet to be purified and its activity tested. However, the sequence of DnpD is most related to the established quinone reductases of TcpB and HadB of the 2,4,6-trichlorophenol degradation pathways in strains JMP134 and DTP0602, respectively (23, 39). Hence, in all likelihood the posing of benzoquinone in the DNP degradation pathway of strain KU-46 leading to the formation of hydroquinone or hydroxyquinol from 4-nitrocatechol is correct as presented in Fig. 1.

For comparison, *Rhodococcus opacus* strain SAO101 degrades NP via a hydroxyquinol pathway whereby a two-component oxygenase/reductase system (*npcA* and *npcB*) is responsible for an initial conversion of NP to 4-nitrocatechol followed by the formation of hydroxy-1,4-benzoquinone and then reduction to hydroxyquinol (40). Interestingly, Yamamoto et al. (43) reported the presence of a hydroquinone pathway in *Rhodococcus* sp. strain PN1 in which the two-component NpsA1A2 hydroxylase system whose amino acid sequences are 100% identical to those of NpcA and NpcB converted NP to 2-hydroxy-1,4-benzoquinone via 1,4-benzoquinone. Previously, strain PN1 was found to contain another two-component hydroxylase system NphA1A2 that converted NP to 4-nitrocatechol in a pathway similar to that of the NpcAB system of strain SAO101 (42). Fig. 9 summarizes the status of NP oxidation and the responsible enzymes that have been cloned from the indicated bacteria. Although not shown in Fig. 9, the 2-chloro-NP degradation system of *Rhodococcus imtechensis* RKJ300 (38) was found to have a *para*-nitrophenol monooxygenase, PnpA1A2, that was identical in amino acid sequences to those of NpcAB, and that they converted 2-chloro-NP to 2-hydroxy-1,4-benzoquinone via chloro-1,4-benzoquinone or chlorohydroquinone to hydroxy-1,4-hydroquinone, indicating that this type of NP monooxygenase had phenol 4-monooxygenae activity. The two component chlorophenol monooxygenases from genera *Burkholderia, Ralstonia* and *Cupriavidus* also exhibit phenol 4-monooxygenae activity (20, 22, 23). Phylogenetic analysis showed that these NP- and chlorophenol monooxygenases, except for *nphA1A2* from *Rhodococcus* sp. strain PN1, belong to the phenol 4-monooxygenase group (Fig. S17). In common they catalyze two sequential oxidations from NP or chlorophenol to hydroxyquinol (48). In contrast, DnpC1C2 catalyzed NP to benzoquinone in a single reaction negating the formation of 2-hydroxy-1,4-benzoquinone unlike the case of NpsA1A2 in strain PN1 (Fig. 9).

**Fig. 9.**
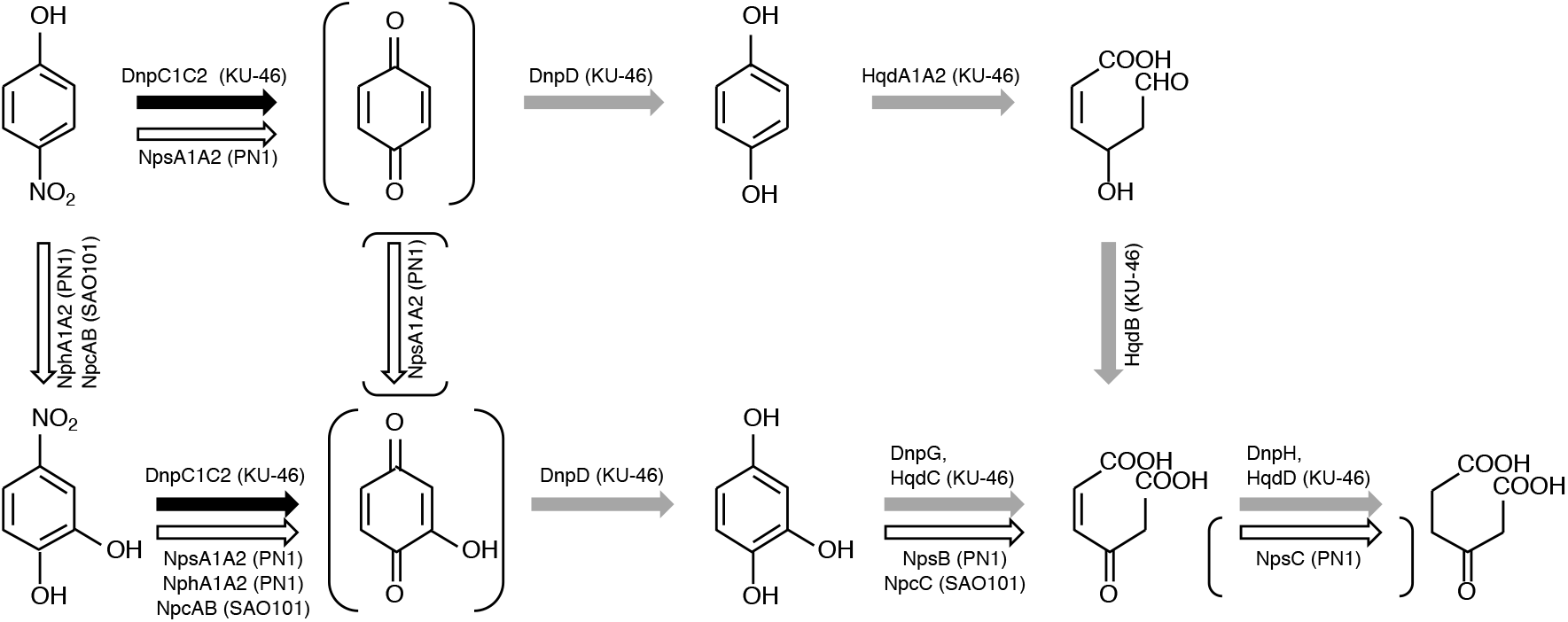
Known 4-NP degradation pathways. Black arrows indicate DpdC1C2 reaction, and gray arrows indicate the reactions presumed with DNA sequence of strain KU-46. Open arrows indicate the pathway from *Rhodococcus* spp., and the two reactions that have not been supported by biochemical results are indicated by the arrows in parentheses.

In the latter part of this study, we isolated a gene cluster *hqdA1A2BDC,* predicted to contain two-component hydroquinone dioxygenase genes, *hqdA1A2*. The *hqdA1A2BDC* genes have the same orientation as the hydroquinone and 4-hydroxyacetophenone degrading genes from *Pseudomonas fluorescens* ACB (49), and the NP degrading genes from *Pseudomonas* spp. and *Burkholderia* sp. strain SJ98 (37, 50–54). In these pathways, the hydroquinone degrading genes are adjacent to the gene encoding nitrophenol monooxygenase and a 4-hydroxyacetophenone monooxygenase, respectively (Fig. S18). *Burkholderia* sp. strain SJ98 has two hydroquinone degrading gene clusters, *pnpABA1CDEFG* and *npcCDEFG* (37). Whereas the *pnpABA1CDEFG* cluster is linked to NP degrading gene, the *npcCDEFG* cluster is not. And because the latter cluster was not upregulated by NP induction Min et al. concluded that the *npcCDEFG* cluster was unlikely involved in NP catabolism (37). In the present case there is no experimental evidence to indicate that the *hqdA1A2BDC* genes in strain KU-46 are directly responsible for NP and DNP degradation. Future studies would attest to this possibility.

In conclusion, this study fills a gap in our knowledge of DNP degradation in a Gram-negative bacterium as well as enhances our understanding of the genetics and biochemical diversity of catabolism of the culpable DNP and derivatives. Further, it reiterates the need to characterize an organism not only to enrich the little we know about microbial diversity at large but also because microorganisms may have new metabolic or biocatalytic properties that can be explored for bioremediation or green processes such as biocatalysis. Lastly, DNP was recently repositioned as a potential disease-modifying drug for a number of insidious diseases in humans such as Huntington, Alzheimer, Parkinson, multiple scelerosis and amyotrophic lateral sclerosis (7). At any rate, knowledge of the many ways DNP can be metabolized will likely be relevant to the possible fate of the drug in human gut microbiota.

## MATERIALS AND METHODS

### Bacterial strains and growth conditions

The bacterial strains and plasmids used in this study are listed in Table 2. *Paraburkholderia* sp. strain KU-46 was grown at 30°C in 1/2 Miller’s LB (Merck Millipore) medium or mineral salt medium (MSM), containing DNP as nitrogen source and succinate as carbon source (15). *E. coli* was grown in Miller’s LB medium or M9 medium containing the following components: 2 mM MgSO_4_.7H_2_O, 0.2 mM CaCl_2_.2H_2_O, 0.002% (w/v) thiamine hydrochloride, 0.002% (w/v) L-proline, and 0.3% (w/v) disodium succinate hexahydrate in M9 salt solution (58). Cultures were incubated at 30°C for the *Paraburkholderia* strains and 37°C for the *E. coli* strains, unless otherwise specified. When necessary, the medium was supplemented with ampicillin (Ap; 100 μg/ml), chloramphenicol (Cm; 20 μg/ml) or kanamycin (Km; 50 μg/ml).

### Isolation and sequencing of DNP degrading gene cluster

Based on the amino acid sequence alignment of the oxygenase components of the two-component phenol monooxygenases (22, 40, 41, 59. 60) two conserved peptide segments (G-N-P-[NED]-H-A-K at position 273-279 in TcpA of *C. pinatubonensis* strain JMP134, and F-E-[NK]-F-N-G-T-P at position 443-450 in TcpA of strain JMP134) were chosen for the design of degenerate primers, TC-FDM-F and TC-FDM-R, (Table 3) for PCR amplification using Hot Start *Taq* DNA Polymerase (New England Biolabs). The amplification conditions were as follows: 95°C for 2 min; 30 cycles at 95°C for 30 s, 55°C for 30 s, and 72°C for 30 s. The 533-bp amplified product was cloned in the vector pXcmkn12 (56) before sequencing, and the resulting plasmid was designated pX-dnpC1 (Table 2).

**TABLE 3.**
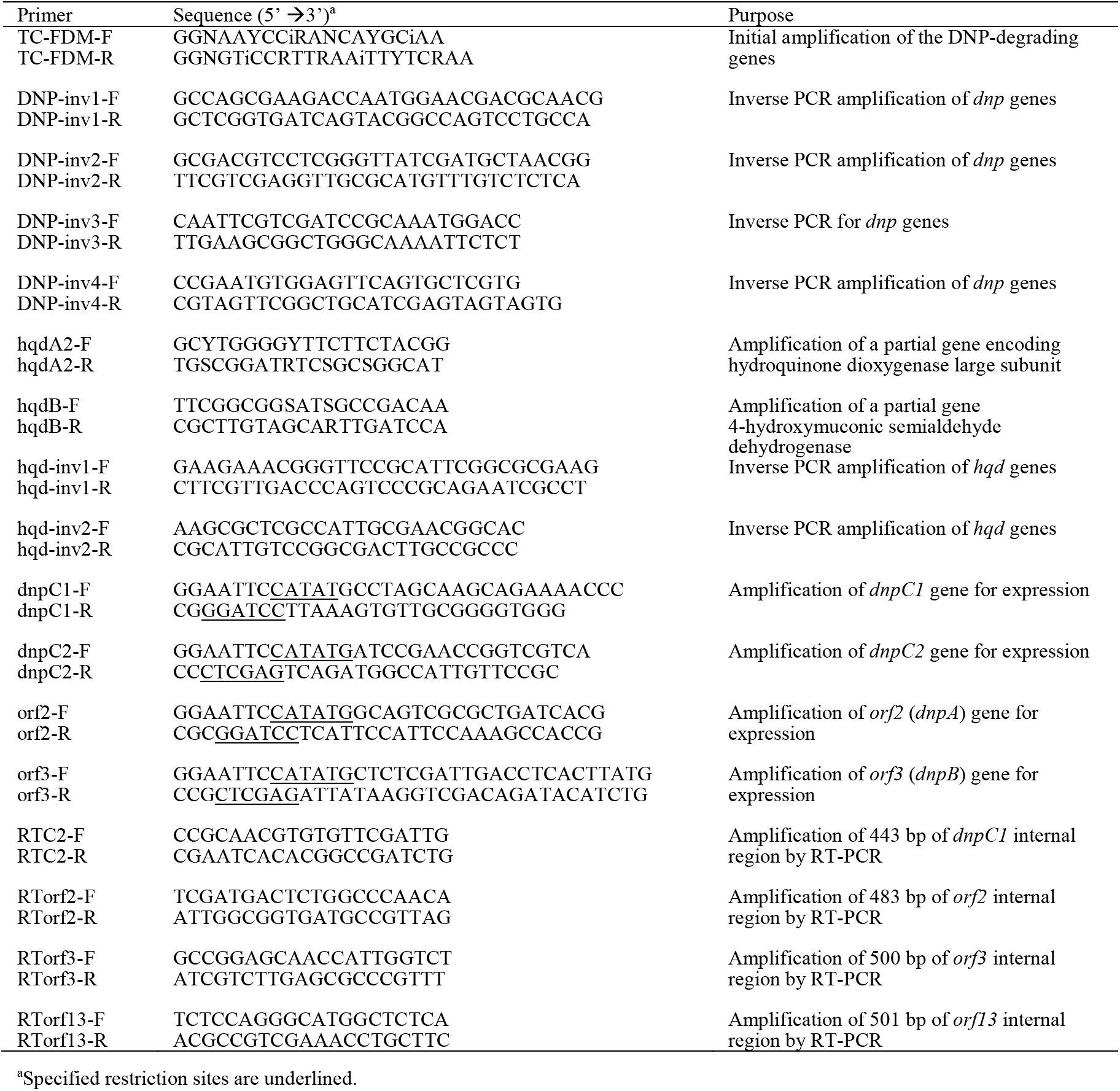
Primers used in this study.

Continuous inverse-PCRs were employed to obtain the flanking fragments with various pairs of primers (Table 3). The inverse PCR was conducted with step-down cycle using KOD FX Neo DNA polymerase (Toyobo) according to manufacturer’s instruction. DNA sequences of the inverse PCR products were determined by direct sequencing and primer walking methods.

The nucleotide sequence was determined with an ABI PRISM 310 Genetic Analyzer using a BigDye Terminator v3.1 Cycle Sequencing Kit (Thermo Fisher Scientific). Analysis of nucleotide sequence and homology searches were performed as described previously (61). Multiple sequence alignment was produced with the program Clustal W (http://clustalw.ddbj.nig.ac.jp).

To obtain the hydroquinone-degradation genes, we first attempted to amplify a portion of the gene encoding the large subunit of the hydroquinone dioxygenase (*hqdA2*) and also a fragment of 4-hydroxymuconic semialdehyde dehydrogenase-encoding gene (*hqdB*). The two sets of degenerate primers, hqdA2-F and hqdA2-R, and hqdB-F and hqdB-R, respectively (Table 3, were designed from the conserved peptide sequences, A-W-G-F-F-Y-G at position 77-83 in HapD of *Pseudomonas fluorescens* strain ACB (49), M-P-A-D-I-R-H at position 299-305 in HapD, F-G-G-[MI]-A-D-K at position 112-118 in HpaE, and W-I-N-C-Y-K-R at position 434-440 in HapE, respectively. PCR amplifications and DNA sequencing were carried out as described above. Inverse-PCRs were employed to obtain the flanking fragments using the primers listed in Table 3.

### Gene disruption

To disrupt the *dnpC1* gene, a 533-bp DNA fragment containing the internal region of *dnpC1* was excised from pX-dnpC1 using BamHI restriction sites of the vector that are adjacent to the inserted fragment, and inserted into vector pK19mob (57). The resulting plasmid, pK19-dnpC1, was introduced by conjugation from *E. coli* S17-1 into strain KU-46. Transformants were selected on MSM plate containing 1.0 g/L ammonium nitrate, 0.3% disodium succinate hexahydrate and kanamycin, and then subjected to PCR analysis to examine insertion of pK19-dnpC1 into the genomic *dnpC1* gene by single crossover (Fig. S1).

### Reverse transcription (RT)-PCR

Cells of *Paraburkholderia* sp. KU-46 were grown in MSM medium containing 0.3% succinate and 0.4 mM DNP, NP or NaNO_3_ until OD_600_ of the culture was 0.3. Cells were then immediately mixed with RNAprotect Bacteria Reagent (QIAGEN). Total RNA was isolated using ISOGEN II (Nippon Gene). In order to remove any contaminating genomic DNA, the RNA samples were incubated with 1 U of deoxyribonuclease (RT Grade) for Heat Stop (Nippon Gene). A cDNA was obtained by a reverse transcription (RT) reaction using PrimeScript II 1st strand cDNA Synthesis Kit (Takara Bio) and random primers. The cDNA was used as a template for subsequent PCRs with specific primers (Table 3). PCR samples were electrophoresed on a 0.8% agarose gel and visualized by staining with ethidium bromide.

### Expression of *dnp* genes in *E. coli*

The DNA fragment carrying *orf2, 3* (*dnpAB*) was amplified by KOD –Plus– Neo DNA polymerase (Toyobo) using the primers shown in Table 3, and the gel-purified DNA was ligated to the expression vector, pET-22b(+) (Novagen). Similarly, the DNA fragments carrying *dnpA*, *dnpB*, *orf4* (*dnpC1*)*, dnpC2,* and *dnpC1C2* were amplified, and ligated to the expression vector, pET16b (Novagen) or pET-HAT. The forward primers were designed to contain an NdeI restriction site with an ATG start codon and the reverse primers contain a EcoRI, XhoI or BamHI restriction site to facilitate directional cloning in the pET-22b(+), pET-16b or pET-HAT expression vectors. The resultant plasmids were designated pET-dnpAB, pET-dnpA, pETHAT-dnpA, pET-dnpB, pET-dnpC1, pET-dnpC2, and pET-dnpC1C2 (Table 2). For protein production, all *E. coli* strains containing expression plasmid including pET-22b(+), pET-16b and pET-HAT as a negative control were cultivated in LB medium containing Ap. When the culture reached an OD_600_ of 0.4-0.5, temperature of rotary shaker was shifted to 25°C, and further cultured to until OD of 0.5 to 0.6 in order to shift the medium temperature to 25°C. At this point (OD_600_ of 0.5 to 0.6), rhamnose was added to induce the protein expression at final concentration of 0.05% in the medium and further cultured for 16 hrs. The resulting cells were harvested by centrifugation, washed twice in 21 mM sodium-potassium phosphate buffer (pH7.1), and used as whole cells.

### Synthesis of H^−^-DNP

H^−^-DNP was chemically synthesized according to the method of Behrend and Heesche-Wagner (29) with minor modifications. 1 mmol of DNP was dissolved in 3 ml of dry acetonitrile and sodium sulfate and heated to 50°C in argon environment for 5 min. At this temperature, mmol of sodium borohydride was added over a period of 20 s. The reaction mixture turned red and further incubated at 50 °C for 3 min. The reaction mixture was cooled on ice, and then an orange red precipitate formed. The supernatant was removed and the precipitate was washed with 3 ml cold argon purge acetonitrile.

As the chemically synthesized H^−^-DNP contained traces of sodium borohydride and other by-products, we used *E. coli* whole cells harboring *orf2* (*dnpA*) to prepare the H^−^-DNP substrate and after cell removal the extract was used for transformation as described below. The molar concentration of H^−^-DNP was considered as the starting material DNP, since the conversion efficiency was 100%. This quantification is consistent in relation to the extinction coefficient value of “ε420=9 mM^-1^cm^-1^ determined by Behrend and Heesche-Wagner method (29).

### Whole cell transformation

The *E. coli* whole cells were resuspended in 21 mM sodium-potassium phosphate buffer (pH 7.1) supplemented with 0.4% glucose, adjusted to an OD of 1.0 at 600 nm and incubated with 0.1 mM of DNP or 0.06 mM of H^−^-DNP. Cell suspensions were shaken at 37 ^°^C, and 5-fold dilution of supernatants were analyzed by a spectrophotometer (UV-1800, Shimazu), and a high-pressure liquid chromatography (HPLC) as described below.

### Protein purification

His_10_-tagged DnpA (H_10_-DnpA), His_10_-tagged DnpB (H_10_-DnpB), and HAT-tagged DnpA (H-DnpA) were purified from *E. coli* cells overproducing the corresponding proteins. The whole cells were resuspended in 21 mM sodium-potassium phosphate buffer (pH7.1), and sonicated by three 40-s bursts with a Braun-Sonifier 250 apparatus. After centrifugation for 30 min at 18,000 X *g* at 4°C, the supernatant was applied to TALON metal affinity resin (Clontech-TAKARA), according to the manufacturer’s instructions. The column containing the crude cell extracts was washed with wash buffer containing 50 mM sodium phosphate, pH7.0, 300 mM NaCl and 20 mM imidazole, and the protein was eluted with an elution buffer containing 50 mM sodium phosphate, pH7.0, 300 mM NaCl, and 120 mM imidazole. Imidazole and NaCl were removed using a PD-10 gel filtration column (GE Healthcare). H_10_-DnpA did not bind to TALON metal affinity resin, therefore we constructed pETHAT-dnpA, which expresses H-DpnA, and purified the H-DpnA protein.

### Enzyme assays

Hydride transferase activity was assayed by monitoring the UV-visible light absorption spectrum change using a spectrophotometer (UV-1800, Shimazu). Reaction mixtures contained DNP or PA (0.04 μmol), NAD(P)H (0.08 μmol), sodium-potassium phosphate buffer (20 μmol, pH7.1), and an appropriate amount of purified enzyme in a final volume 1 ml. Nitrite-eliminating enzyme was also assayed by monitoring the UV-visible light absorption spectrum change using a spectrophotometer (UV-1800, Shimazu). Reaction mixtures contained H^−^-DNP (0.05 μmol) sodium-potassium phosphate buffer (20 μmol, pH7.1), and an appropriate amount of purified enzyme in a final volume 1 ml.

### HPLC analysis

HPLC analysis was performed on a CAPCELL PAK C18UG120 column (column size of 4.6 by 250 mm and particle size of 5 μm; Shiseido) connected to LC-6AD pump and a SPD-M20A photodiode array detector (Shimadzu). For the analysis of transformation of NP, the mobile phase consisted of methanol/H_2_O, 1:1 v/v, containing 0.07% perchloric acid at a flow rate 1.0 ml min^-1^. For the analysis of transformation of DNP and H^−^-DNP, the mobile phase consisted of 50 mM potassium phosphate (pH 8.0)/methanol, 3:2 v/v, at a flow rate of 1.0 ml min^-1^.

### Nucleotide sequence and accession numbers

The DNA sequences of the gene clusters *dnp* (14.8 kb) and *hqd* (6.8 kb) had been deposited in the DDBJ database under accession numbers LC496529 and LC496530, respectively.

## Acknowledgments

We thank Dr. Takaaki Sumiyoshi for synthesis of H^−^-DNP. This work was supported in part by a Grant-in-Aid from Kansai University for progress of research in a graduate course, 2018.

